# Mycelium Biocomposites from Agricultural and Paper waste: Sustainable alternative to plastic foam based secondary packaging

**DOI:** 10.1101/2025.02.18.638821

**Authors:** Sandra Rose Biby, Vivek Surendran, Lakshminath Kundanati

## Abstract

Plastic foams, which were once lauded as a breakthrough invention in packaging and insulation industries, have gradually evolved into a global problem and are more concerning in developing nations because of ineffective waste management systems. The convenience and pervasiveness of these come at a high cost with adverse effects on human health and our planet’s ecosystems as they are not biodegradable. It is pivotal that an alternative and sustainable solution is found to address this problem. Mycelium-based biocomposites, made up of root-like networks called hyphae are sought out as a potential substitute for plastic-based foams as they are biodegradable. They employ a waste-to-value strategy by using agricultural wastes as a growing media for the mycelium. The study primarily focuses on the comparative analysis of mycelium-based biocomposites with existing foam materials such as Expanded PolyStyrene (EPS) and evaluates their efficacy as a biodegradable packaging material. Thus, we plan to identify the efficacious strain of mycelium from *Pleurotus ostreatus* or *Ganoderma lucidum* and also explore various substrate materials such as cocopith, cardboard, paper, and sawdust. Our preliminary compression results show that the biocomposites can be engineered to have Young’s modulus ranging from 1-3 MPa for both strains. These results show that mycelium-based biocomposites can be a promising alternative to EPS /Styrofoam and move towards a sustainable solution that can reduce the dependence on non-biodegradable foam-based materials in developing nations like India.

## 1. Introduction

Plastic, which was once lauded as a breakthrough invention, has gradually evolved into a global problem. Plastic, as a flexible and long-lasting material, has made its way into nearly every facet of our lives, from packaging and building to healthcare and technology. However, the convenience and pervasiveness of plastic come at a high cost, with negative effects for our planet, ecosystems and eventually our own health (Kumar et al).The amount of plastic waste produced globally is enormous. According to the Allen McArthur Foundation, just 9% of all plastic garbage generated is recycled, 12% is burnt, and 79% is dumped into landfills after its useful lives(Geyer, Jambeck and Law, 2017; S Walker and Rothman, 2020; Shams, Alam and Mahbub, 2021; Kibria et al., 2023). CPCB reports says that 4,126,997 TPA of plastic waste was generated in the year 2020-21. The widespread usage of plastic has resulted in a serious environmental disaster. Plastics are primarily sourced from fossil fuels, thus depleting these limited resources. Furthermore, the manufacturing process emits significant volumes of greenhouse gases into the environment, aggravating climate change. Once dumped, plastic trash has a devastating impact on our ecosystems. Non-biodegradable plastics clog landfills, contaminating soil and water supplies, while massive volumes of plastic trash collect in our oceans, harming marine life and destroying entire ecosystems (S Walker and Rothman, 2020).Given these significant concerns, it is critical that we address the negative effects of plastic usage and move towards long-term solutions. We can reduce the environmental, health, and economic difficulties associated with plastic pollution by lowering our reliance on single-use plastics, increasing recycling and waste management, and stimulating the development of alternative materials. Recycling has emerged as an important strategy for reducing the impact of plastic waste. It is a promising option since it redirects plastic materials away from landfills and incinerators, lowering the demand for plastic manufacture, and conserving important resources. We can contribute to a cleaner, more sustainable future by embracing recycling practises on both an individual and societal level (Okunola A et al., 2019).

Recycling can be an effective way to cut down on plastic waste, but it can also be a major source of microplastic production. Microplastics are formed mostly through the fragmentation and disintegration of larger plastic products, such as plastic trash in the environment, as well as the breakdown of synthetic materials with dimensions ranging from 1µmto 5 mm. They are minute plastic grains that are commonly found in abandoned plastic fragment goods. They are carried to the sea by rivers and floods, and they often choke canals, causing pollution and are potential risk to human health(Campanale *et al*., 2020a, 2020b; Bhuyan, 2022a, 2022b; Brown *et al*., 2023a, 2023b).As a result, recycling is not always the best solution, therefore a biodegradable and compatible alternative option must be developed (Wojnowska-Baryła, Bernat and Zaborowska, 2022).In contrast to the conventional linear economic model, which follows the ‘take-make-consume-throw away’ pattern, a circular economy focuses on products and the materials they include, and is built on reuse, repair, refurbishing, and recycling in an almost complete loop. It means cutting waste to the absolute lowest(Elisha; Meyer et al., 2020; Kirchherr et al., 2023). Mycelium-based biocomposites adopts a circular economy strategy in which the substrates used are agricultural waste byproducts such as sawdust, coconut coir etc., which are frequently discarded as waste material. These materials can be reused with great efficiency and are totally decomposable. They also complete the loop of the circular economy by utilizing these as substrates for mycelium-based byproducts.

Algal-based bioplastics are a growing alternative for biodegradable packaging that can effectively replace commercial plastic in the direction of a sustainable circular economy. They are a viable resource for the long-term, sustainable manufacturing of bio-based materials in large quantities(Yap *et al*., 2023). The pseudo stem of bananas has a high fibre content (∼5%) and is currently underutilized. Instead of using styrofoam sleeving, this biodegradable raw material is being used to create fruit wrap paper from leftover banana pseudo-stems(Yoga Milani *et al*., 2020).Leather can also be made from pineapple leaves. The material is comprised of fibre from pineapple leaves and is natural and non-woven. The leaves are a byproduct of the present agriculture and fruit sector, and they would otherwise be discarded. This material is cruelty-free, naturally sourced, and sustainably sourced(Duangsuwan et al., 2023).

Fungi are one of nature’s most astonishing biodegradable champs. These intriguing organisms have unique features like rapid growth, adhesiveness, biodegradability, versatile substrate compatibility make them ideal for decomposing complex organic compounds(Frąc, Silja E. Hannula, et al., 2018). Fungi, such as particular types of mushrooms and moulds, hold significant potential for long-term waste management and environmental preservation. Fungi excel at biodegradation because of their ability to create a varied array of enzymes. Enzymes function as biological catalysts, allowing complex compounds to be broken down into smaller components. Fungi secrete a variety of enzymes, including cellulases, ligninases, and lipases, which are capable of efficiently breaking down chemical bonds in various substrates.

Mycelium materials, a new substance made up of filamentous fungi and agricultural substrates, have the potential to be a valuable by-product. Mycelium feeds on substrates rich in cellulose or hemicellulose, such as rice straw, banana leaves, coffee grounds, coconut coir, or other agricultural wastes. Fungi can be inculcated onto these substrates to form mycelium-based biomaterial once the substrate is processed and provided with essential growing conditions. Mycelium grows by apical tip expansion and the use of hyphae from a spore or an inoculum. Following a period, the hypha begins to branch and form tree-like colonies. They interact, forming a network-like structure and eventually functions as an adhesive. They also expand outwards, forming a dense white fungal skin coating that protects and reinforces the composite (Aiduang et al., 2022b; Tunlid et al., 2022; Balaeș, Radu and Tănase, 2023b).

Mycelium bio composite is not only biodegradable, but it is also a method of recycling agricultural waste. According to Indian council of agricultural research tonnes of agricultural waste are generated worldwide each year, with India creating about 350 million ton. Agricultural wastes or crop residues are either burned or left in fields to decay; nevertheless, burning can emit hazardous pollutants into the atmosphere but they have the potential to be good substrates. Considering the wide availability of these agricultural wastes in India, they can be used to make mycelium-based bio composites which can contribute to reduction in foam-based packaging (Kundanati, 2022). All cellulose-based agricultural wastes, such as sawdust, cardboard, and paper wastes and combinations of two or more substrates, can serve as ideal substrates for mycelium-based biocomposite. Using agricultural waste as a substrate for producing mycelium-based bio composites can thus be a cost-effective method of utilising agricultural waste while lowering waste output and safeguarding the environment(Duque-Acevedo et al., 2020; Koul, Yakoob and Shah, 2022).

*Pleurotus* species, such as *Pleurotus ostreatus* (oyster mushroom), have been studied for the capacity of their mycelium to adhere and grow on a variety of substrates like straw, hemp. *Pleurotus* mycelium has good adhesion qualities, which aid in the creation of robust and cohesive biocomposites(Hoa and Wang, 2015; Xing et al., 2018).*Trametes versicolor*, known as turkey tail, has gained popularity as a component of mycelium-based biocomposites. *Trametes* mycelium has shown potential fire-resistance qualities, making it appropriate for applications requiring flame retardancy(Jones *et al*., 2020)*.Ganoderma* species such as *Ganoderma lucidum* and *Ganoderma applanatum* have been investigated for their ability to produce mycelium-based biocomposites. These strains grow quickly and have promising mechanical qualities, making them appropriate for applications that require strength and durability(Obire; Patricia and Cabrera, 2018). The main goals of the study are to compare various mycelium-based bio composites, identify the best substrates for mycelium growth (cocopith, sawdust, paper, and cardboard), evaluate how effective these substrates are as biodegradable packaging materials, and determine which strain of *Pleurotus ostreatus* and *Ganoderma lucidum* is the most effective. Furthermore, the composite’s density, strength, water absorption, were examined.

## 2. Materials and methodology

### 2.1 Inoculation and moulding

Materials including cardboard, paper, sawdust and cocopith were used as substrates. The substrates were chopped and shredded into fine pieces. They were then soaked in distilled water for 24 hours and sterilized (autoclaved at 121°C for 20 minutes) before use. *Pleurotus ostreatus* and *Ganoderma Lingzhi* was purchased from Nuvedo labs, Bangalore. The substrates were then combined with a fixed quantity of fungal inoculum (1:1). The mixed material was then uniformly distributed into sterile moulds of desired shape and size.

### 2.2 Incubation and drying

The sterile moulds with substrates and inoculum were incubated for 2 weeks under standard ambient conditions at 25-30 degrees Celsius. Peptone water is prepared and sprayed by combining 0.5g of peptone with 100ml of distilled water. Peptone water includes a variety of nutrients that stimulate the growth of *Pleurotus ostreatus* and *Ganoderma lingzhi*. It contains amino acids, peptides, nitrogen compounds, vitamins, minerals, and polysaccharides that promote mycelium growth. After incubation the specimens were dehydrated, and the fungal growth was terminated by keeping them in a hot air oven set at 50 °C for 48 hour.

### 2.3 SEM Analysis

A small piece of 2 x 2 cm length of the outer layer of prepared mycelium samples were carefully cut out and mounted on double-sided carbon tape, attached to an aluminium stub, and placed in a desiccator to avoid moisture and dust accumulation. As sample were nonconductive, they were coated with a thin layer of gold with sputter coater (QUORUM, SC7620, made in UK). A scanning electron microscope (Zeiss,evo 18, Germany) was used to image the samples in different magnification.

### 2.4 Optical imaging

Images of growing biocomposites samples were taken and observed using stereomicroscope (Euromex – NexiusZoom EVO, Netherlands) to monitor the changes in the top layer and outer surface appearance of the biocomposites.

### 2.5 Compression test

Cylindrical samples were sectioned out from the moulded samples using laser cutting and we have tested 2 samples each from each set of the samples prepared from each combination of substrate and spawn. Compression strength was determined using a Shimadzu EZ-LX, load bench with a 5 kN capacity and a 500N load cell under ambient conditions (25 °C with 40 to 50% relative humidity). The tests were conducted with controlled displacement at a rate of 1 mm/min. The load–displacement curve was converted to a stress–strain curve using the following formulas to calculate the compressive stress σ and the strain ε: Stress σ = F/A and Stress ε = ΔL/L_o_, respectively (where F: compressive force (N), A: original cross section of the specimen (mm^2^), ΔL: displacement of the loading surfaces (mm), and L_o_: original height of the test piece (mm)).

### 2.6 Water absorption tests

Disk-shaped specimens of 8cm diameter were tested to measure their water uptake when placed on top of water. Specimens were placed in containers filled with distilled water maintained at 30 ± 1 °. Each material’s water absorption rate was measured at zeroth hour, 1 hour, 2 hours, 3 hours, 4 hours, 24 hours, and 48 hours. The weight of the materials and their percentage increase in weight over time were recorded. For each measurement, samples were manually retrieved from the water and weighed using a weighing machine within 1 minute. The collected data was used to calculate the water absorption percentage using the following equation

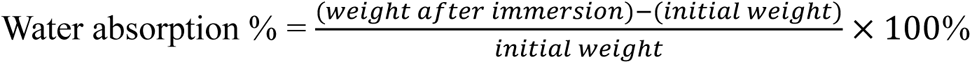

### 2.7 Water contact angle test

The water contact angle of grown bio-composites was determined using a goniometer. The measurements were made on 2 locations of each of the samples. The sample was secured to the instrument platform and a 2µl drop of distilled water was dropped to the material’s surface using a syringe. The image of the droplet was captured using OneAttension by Biolin scientific software, and the water contact angle was calculated. Each sample was tested in duplicates. Finally, an average value was taken. A contact angle less than 90° indicates that the material is hydrophilic or poorly water resistant, whereas a contact angle more than 90° indicates that the substance is hydrophobic or highly water resistant.

### 2.8 Biodegradation test

The mycelium based biocomposites were placed in a nylon mesh bag and then buried in potting soil for different time interval (Van Wylick *et al*., 2022a). The changes in the samples appearance and weight loss percentage were monitored. The samples were taken out every week and the soil were dusted using a brush, dried in hot air oven at 45°C for 3-4 hrs to remove the moisture. A relative humidity of 60-70% was maintained. The data was gathered, and the following equation was used to determine the percentage of weight loss:

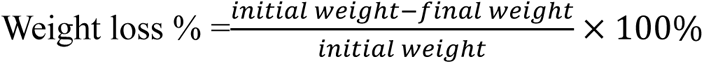

## 3. Results and discussion

### 3.1 Optical imaging

The visual appearance of all samples became full white, the whole surface of the sample was covered with mycelium making it look fully white in colour on day 15 as in **Figure. 1**, indicating that *Ganoderma lucidum* and *Pleurotus ostreatus* mycelium had grown from substrates. The process of creating mycelial composite involved the fungus breaking down substrates through the secretion of cellulase, lignin peroxidase, and laccase to release vegetative growth substances. An earlier study highlighted that the growth *of Ganoderma lucidum* mycelium was denser on sawdust compared to *Pleurotus ostreatus*, which exhibited sparser growth on cocopith and hay (Peng *et al*., 2023). This suggests that different substrates influence mycelial density and overall growth, with sawdust being more favourable for

**Figure 1.** Optical image of samples after 15 days of growth.

*Ganoderma lucidum* due to its ability to secrete enzyme that break down the substrate effectively. This resulted in the growing mycelium and substrates becoming entangled and bonded to one another, strengthening the natural fiber or bonding the core material. All the samples had dense mycelium covering their outer surfaces, although each sample’s surface mycelial density varied. In comparison, the mycelial layer on the outer surface of hay samples were comparatively sparse, and some substrate fibers is still visible. This indicated that hay substrates were less favourable to mycelial growth. The fibrous nature of hay and loose pores in cocopith can create a less cohesive structure for mycelium to colonize compared to denser substrates like sawdust or cocopith. This can lead to uneven colonization and reduced binding efficiency within the composite material. The loose structure may also make it more challenging for the mycelium to establish a strong network necessary for binding the substrate particles effectively. It was also found that sawdust substrate of *Ganoderma lucidum* sample produced denser mycelium layer when compared to *Pleurotus ostreatus*(Elsacker, Vandelook, Brancart, Peeters and Laet, 2019; Nashiruddin *et al*., 2022).

### 3.2 SEM and EDAX Analysis

SEM pictures of the surfaces and interiors of mycelium bio-composites formed from various substrates are presented in **Figure 2**. Several variations in the mycelium microstructure were detected. A layer of mycelial film coated the external surfaces of all the materials; however, the materials differed in terms of surface porosity and roughness. The hypha’s diameter was primarily found within the 1.4–4.0 µm range (measured using image J software). The surface of the *Ganoderma lucidum* strain sawdust biocomposite was less rough and had less pores. The sawdust sample had the densest hyphae, with diameter ranging between 3.01 and 4 µm hyphae. The degradation of lignin plays a crucial role in the structural integrity of the samples. Laccase and peroxidase enzymes present in sawdust facilitate in lignin breakdown which leads to more compact mycelial network(Wang et al., 2024). In contrast, the hyphae of the *Pleurotus ostreatus* strain’s sawdust biocomposites ranged in diameter from 1.5 to 1.9 µm(Haneef *et al*., 2017).

**Figure 2.**
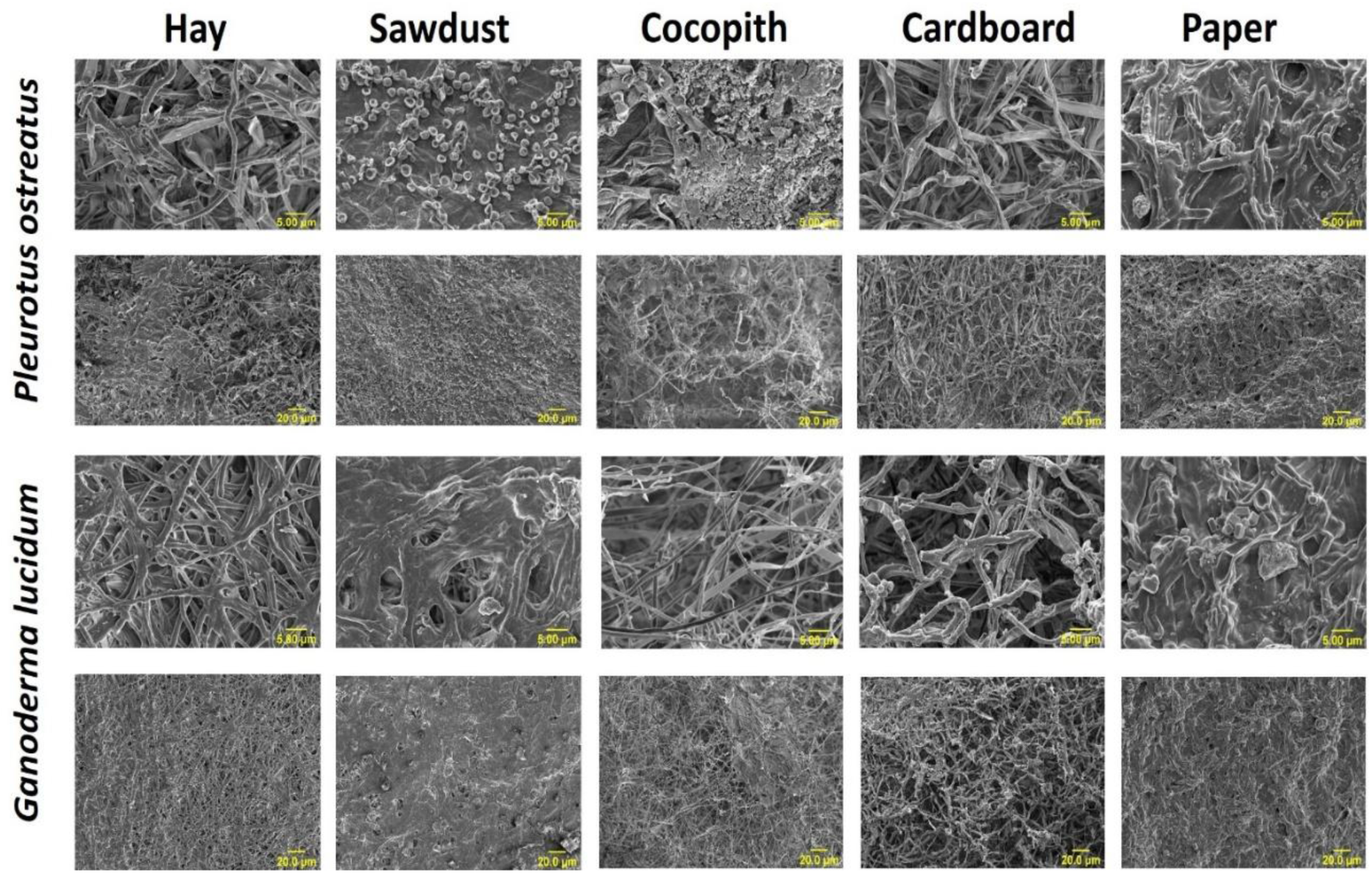
SEM images of *Pleurotus ostreatus* and *Ganoderma lucidum* samples at scale bars of 5 and 20 µm.

The hypha in *Ganoderma lucidum* hay samples developed in a planar network and tended to stick collectively. The diameter of the hyphae varied between 1.5 and 2.01 µm. Paper and sawdust samples of *Ganoderma lucidum* and *Pleurotus ostreatus*, respectively, showed the presence of oxalate crystals as seen in **Figure 3**. These crystals are generated by fungi and have been described in the literature as electron donors in the breakdown of lignocellulose. They are discharged outside the fungal cell, as observed in the SEM images(Frank-Kamenetskaya *et al*., 2021). The hyphae of *Pleurotus ostreatus* strain of cardboard biocomposites ranged in diameter from 2-2.14 µm whereas the hyphae of *Ganoderma lucidum* strain of cardboard biocomposites ranged in a diameter from 2.4-2.7 µm. Among the biocomposites made from *Pleurotus ostreatus* samples, the hyphae of cocopith samples were dense and had a diameter ranging from 2.7-3.09 µm.

**Figure 3.**
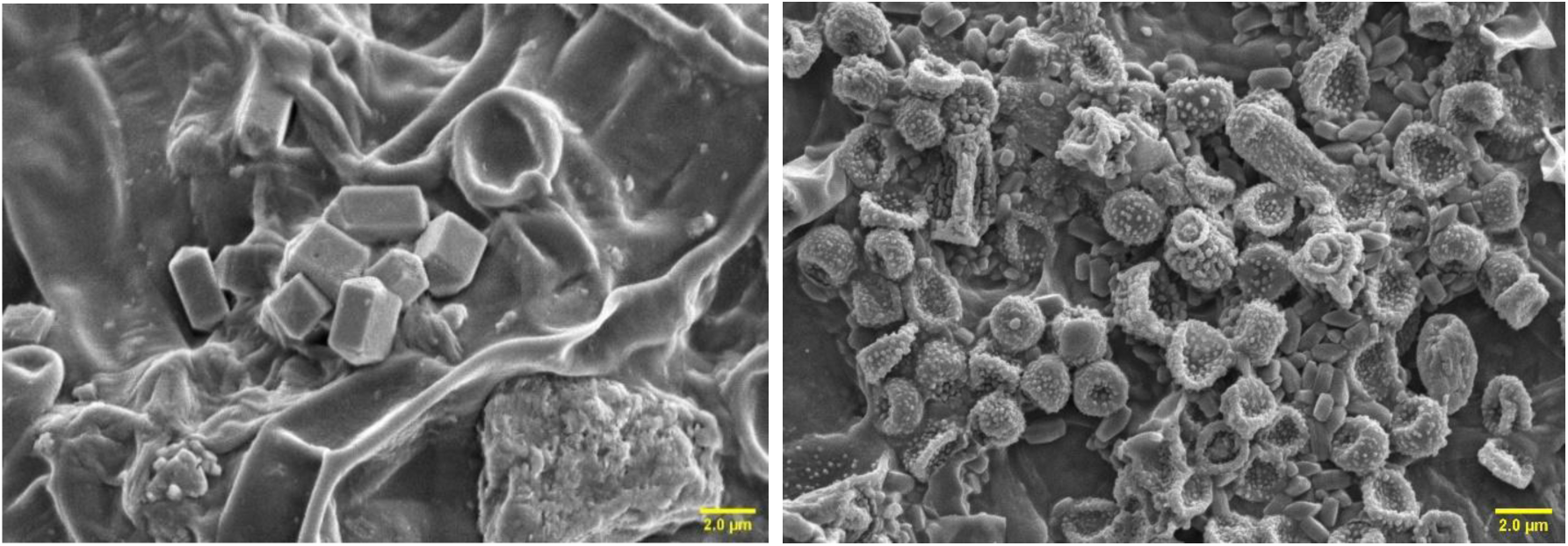
SEM image of oxalate crystals on paper and sawdust samples of *Ganoderma lucidum* and *Pleurotus ostreatus*, respectively samples respectively

Significant peaks in the EDAX graph at approximately 4.0 keV and 3.7 keV indicate the presence of calcium **(Figure 4.)**. A peak near 0.3 keV suggests the presence of carbon. Additionally, a peak around 3.3 keV points to the presence of potassium. Presence of calcium and carbon is a potential indicator of oxalate crystals. The formation of these crystals typically occurs when oxalic acid, secreted by the fungi, interacts with calcium ions present in their growth substrates. This process not only helps in nutrient cycling but also enhances the structural properties of the mycelium *(Xiao* et al.*, 2022)*.

**Figure 4.**
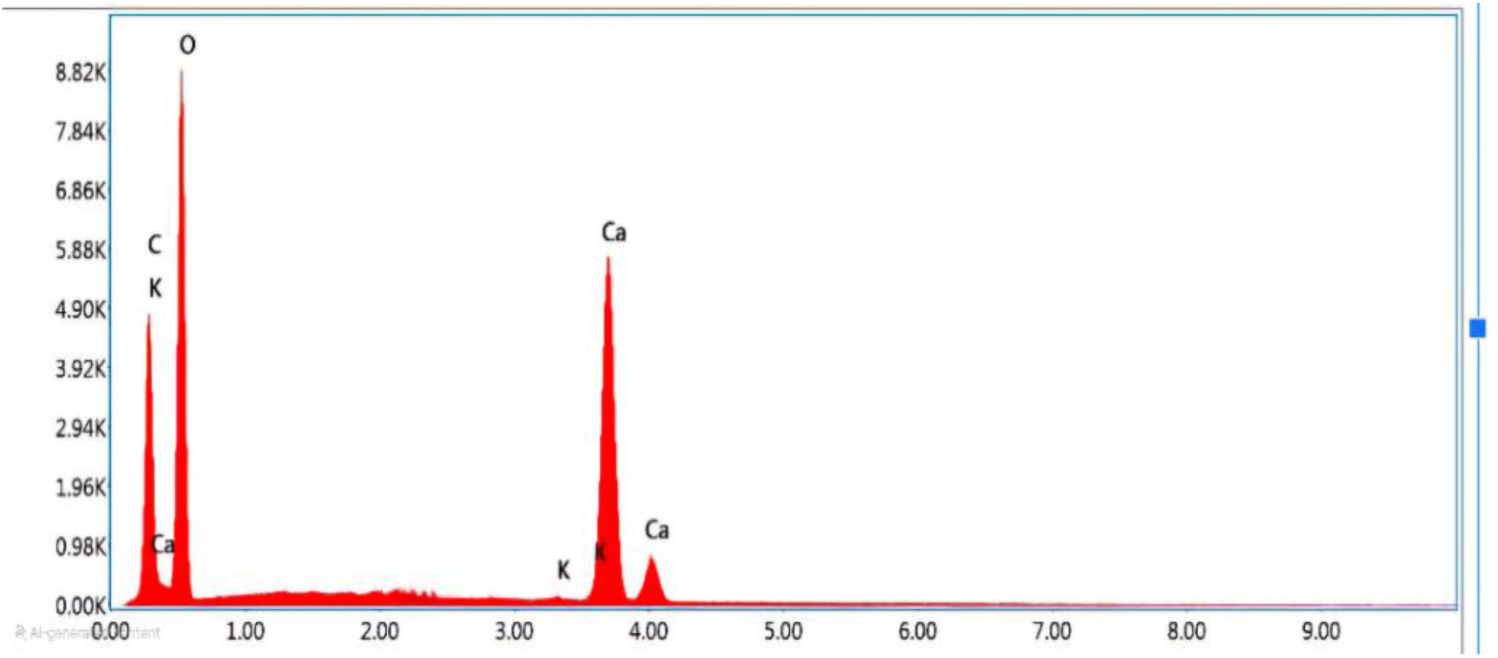
EDAX graph of oxalate crystals

### 3.3 Compression test

The compression test results of various substrates showed notable differences in their mechanical properties. The compression strength of the biocomposite was highly related to the type of substrates. The accumulation of chitin from the fungal cell wall in the substrate material particles could provide the material a mechanical resistance similar to that of EPE and EPS and lessen material cracking under compression(Peng *et al*., 2023). Among the susbtsrtaes used, cardboard demonstrated the highest compressive strength as depicted in **Figure 6**. for *Ganoderma* at 2.72±0.41 Mpa, exceeding *Pleurotus*, which measured only 0.83±0.07 Mpa. In cocopith *Ganoderma* exhibited higher strength at 1.79± 0.72MPa compared to *Pleurotus* 0.91±0.10 MPa. In contrast *Pleurotus* sample of sawdust substrate showed better compressive strength of 1.79± 0.02 MPa when compared to *Ganoderma* which had a strength of 1.52± 0.54. Similar results were observed by (Yang *et al*., 2017), indicating that sawdust substartes exhibits superior compressive strength. The thick chitin layer formation around the substrates formation in paper substrates can be a reason for its high compressive strength (Dutton and Evans, 1996). Hay exhibited lowest compressive strength for both strains and also a lot of variation primarily due to heterogenous compaction during substrate seeding that is dependant on the particle length. Also, the composites had high porosity that resulted in less strucutrally integral leading to samples fracturing early in the compression tests.

In comparison to EPE foam sample as in **Figure 5**, had an elastic modulus of 0.69±0.006 Mpa which is significantly lower than the modulus values observed for *Ganoderma* on cardboard and cocopith but higher than that of *Pleurotus* on cardboard and hay substrate. Thus, EPE foam is less stiff compared to the biocomposites derived from *Ganoderma* on specific substrates(Sivaprasad et al., 2021). On the otherhand, EPS foams exhibited a modulus value of 0.281 ±0.05 MPa. Thus, the biocomposites need fruther tuning of substrate compostion to match the elastic properties of the EPS, which is one of the widely used secondary packaging material. The overall findings indicate that *Ganoderma* generally exhibits higher compressive strength on most substrates, *Pleurotus* excels on sawdust highlighting the importance of substrate selection in optimizing growth conditions for each fungal strain(Tacer-Caba *et al*., 2020). This can be mainly due to the higher hyphae thickness of *Ganoderma* strain compared to *Pluerotus.*Thicker hyphae can provide greater structural support, contributing to higher compressive strength in certain substrates.

**Figure 5.**
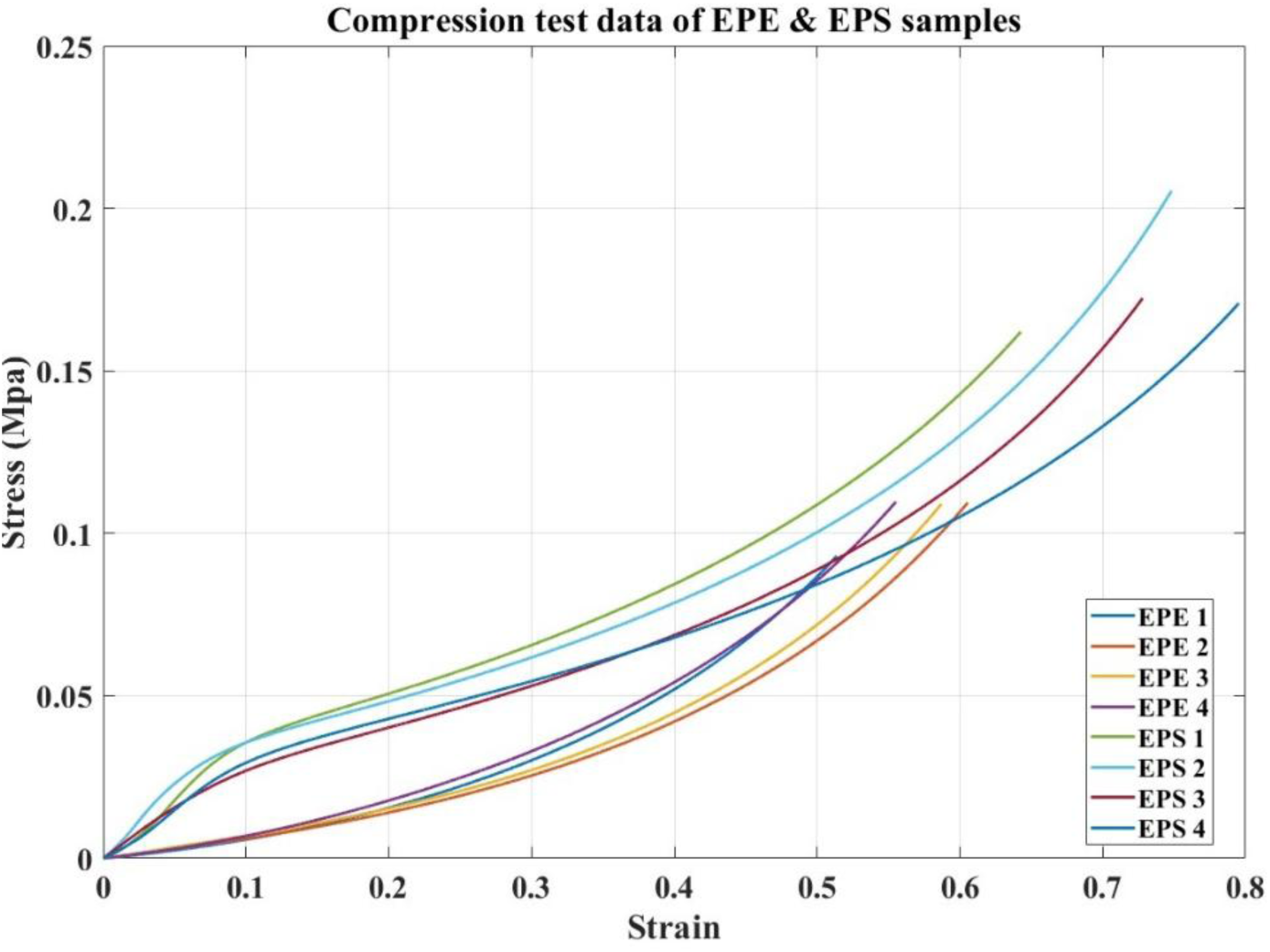
Compression test graph of EPE and EPS foam samples

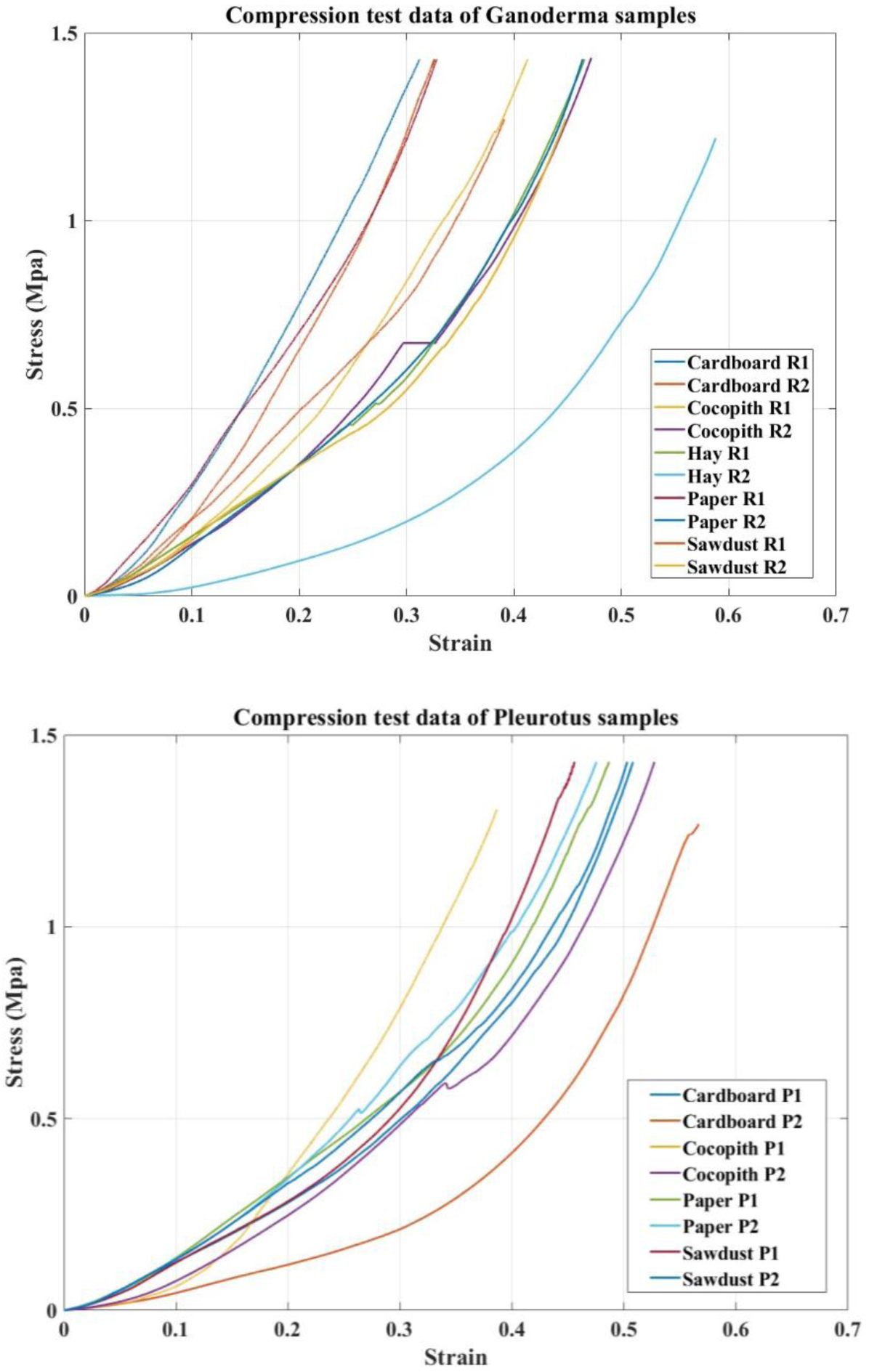

**Table 1.**
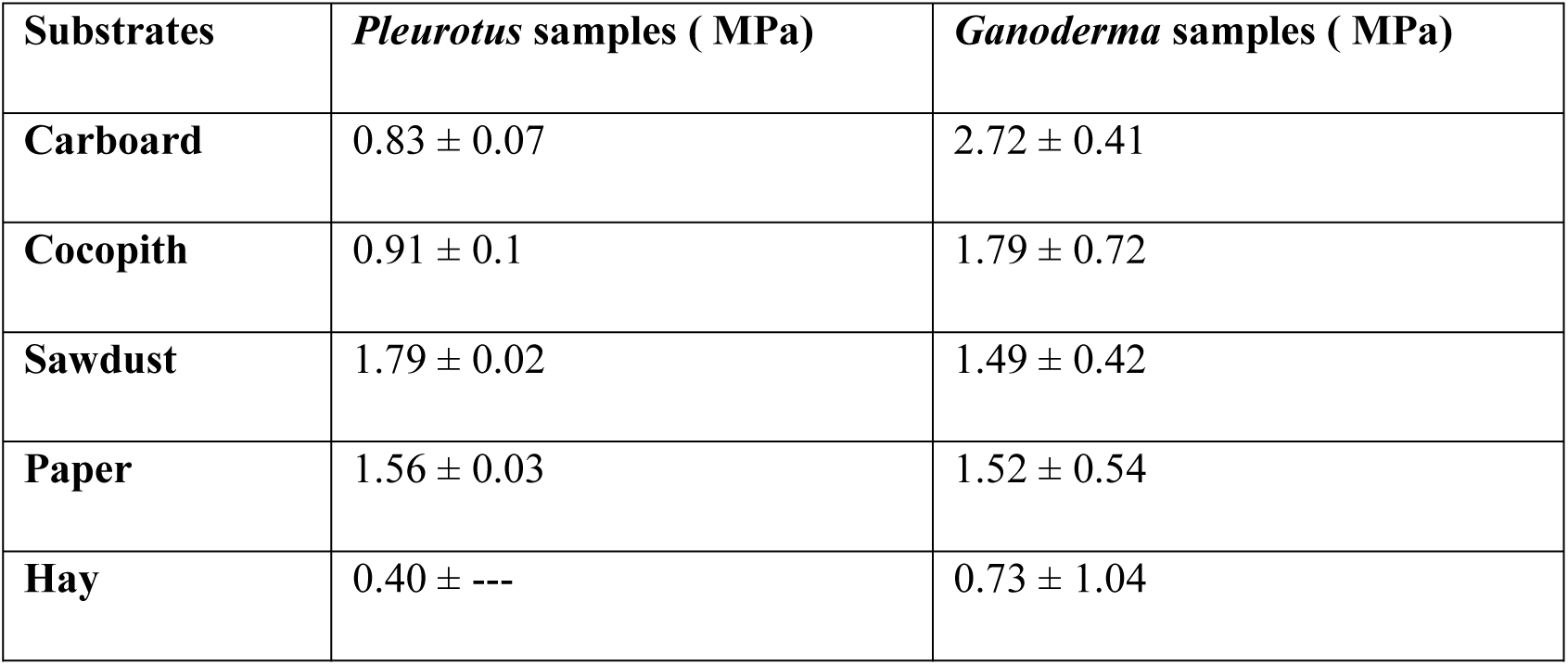
Elastic modulus values calculated from the compression tests.

**Table 2.** Elastic modulus of EPE and EPS samples.

| Elastic modulus of EPE samples in MPa |  |  |
| --- | --- | --- |
| Samples | E (MPa) | Average E value |
| Sample 1 | 0.066 | 0.069 ±0.006 |
| Sample 2 | 0.063 |  |
| Sample 3 | 0.069 |  |
| Sample 4 | 0.078 |  |
| Elastic modulus of EPS samples in Mpa |  |  |
| Samples | E (MPa) | Average E |
| Sample 1 | 0.356 | 0.281 ±0.05 |
| Sample 2 | 0.251 |  |
| Sample 3 | 0.227 |  |
| Sample 4 | 0.292 |  |

### 3.4 Water absorption test

Water absorption rates of different biocomposites over various time intervals demonstrated different trends (**Figure. 7**). The results show that all materials exhibited an increase in weight over time, with sawdust (increased by 462 % in 48 hrs) demonstrating the highest percentage increase, followed by cardboard (409%), paper (327.8%), cocopith (210%) and hay (175%) of *Ganoderma linghzhi* samples. Hay biocomposites made of *Ganoderma linghzhi* strain absorbed less water. In contrast hay biocomposites of *Pleurotus ostreatus* samples absorbed higher percentage of water (415.5%) followed by paper (286.1%), sawdust (263.5%), cardboard (183%) and cocopith (171%) in 48hrs.

**Figure 7.**
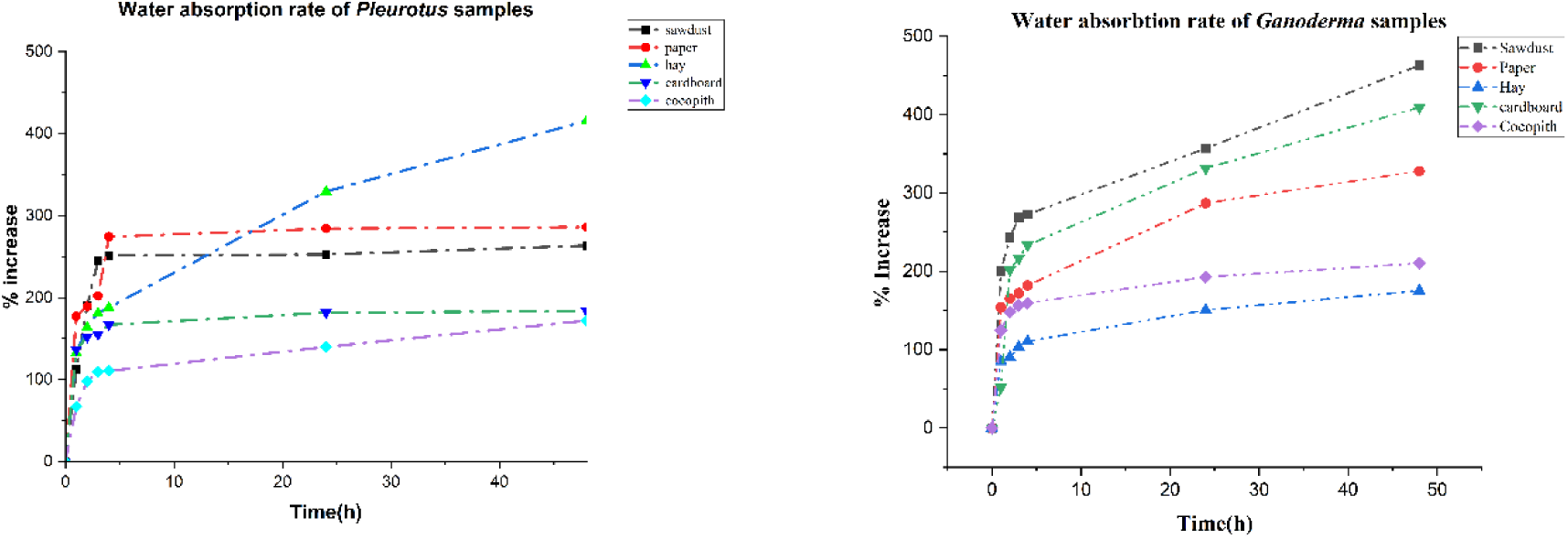
Water absorption rate of *Pleurotus ostreatus* samples and *Ganoderma lucidum* samples.

**Figure 8.**
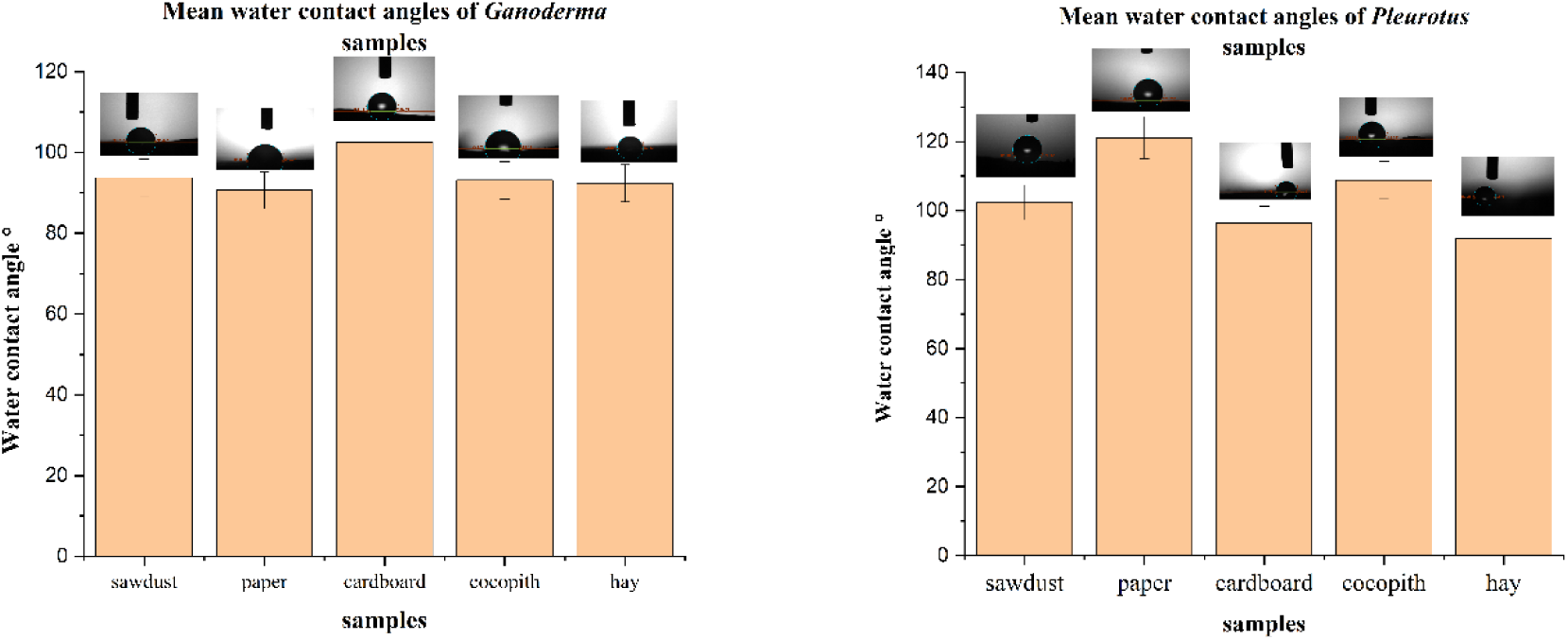
Water contact angle measurements of *Ganoderma* and *Pleurotus* samples

This can be mainly due to the higher density and interconnection of hyphae of *Ganoderma lucidum* samples where the diameter of hyphae ranged between 2.01-4.00 µm when compared to *Pleurotus ostreatus* ranging from 1.00-3.09µm(Elsacker et al. 2019; Lee and Choi, 2021).

### 3.5 Water contact angle test

The surface water contact angle (WCA) of mycelium bio-composites measured from various substrates is shown below in **Figure 9**. The average WCA of the sawdust substrate in the *Pleurotus* sample was 121.98°, and the WCA of the cardboard substrate in the *Ganoderma* sample was 109.72°. Other materials had a WCA greater than 90°, which was higher than the water contact angle of the EPS material (90°). This supports the use of these biocomposites for use of packaging that encounter water for short spans of time. It is reported that the exterior layer of pure mycelial material was hydrophobic, because hydrophobin, a self-assembling protein, blanketed the aerial mycelium’s surface(Evgeniya Lock, 2008; Antinori et al., 2020). The water contact angle of the sawdust substrate in the *Pleurotus* sample and the cardboard substrate in the *Ganoderma* sample were slightly greater, indicating that their surfaces are more hydrophobic. It was primarily caused by the growth of a full mycelial biofilm on its surface and the lack of holes. In addition, the water contact angle of mycelium bio-composite was proportional to the density of mycelium growth. Denser mycelium growth and a flatter surface with fewer pores were favorable for forming a greater WCA of material (Yuan and Lee, 2013; Peng et al., 2023).

**Figure 9.**
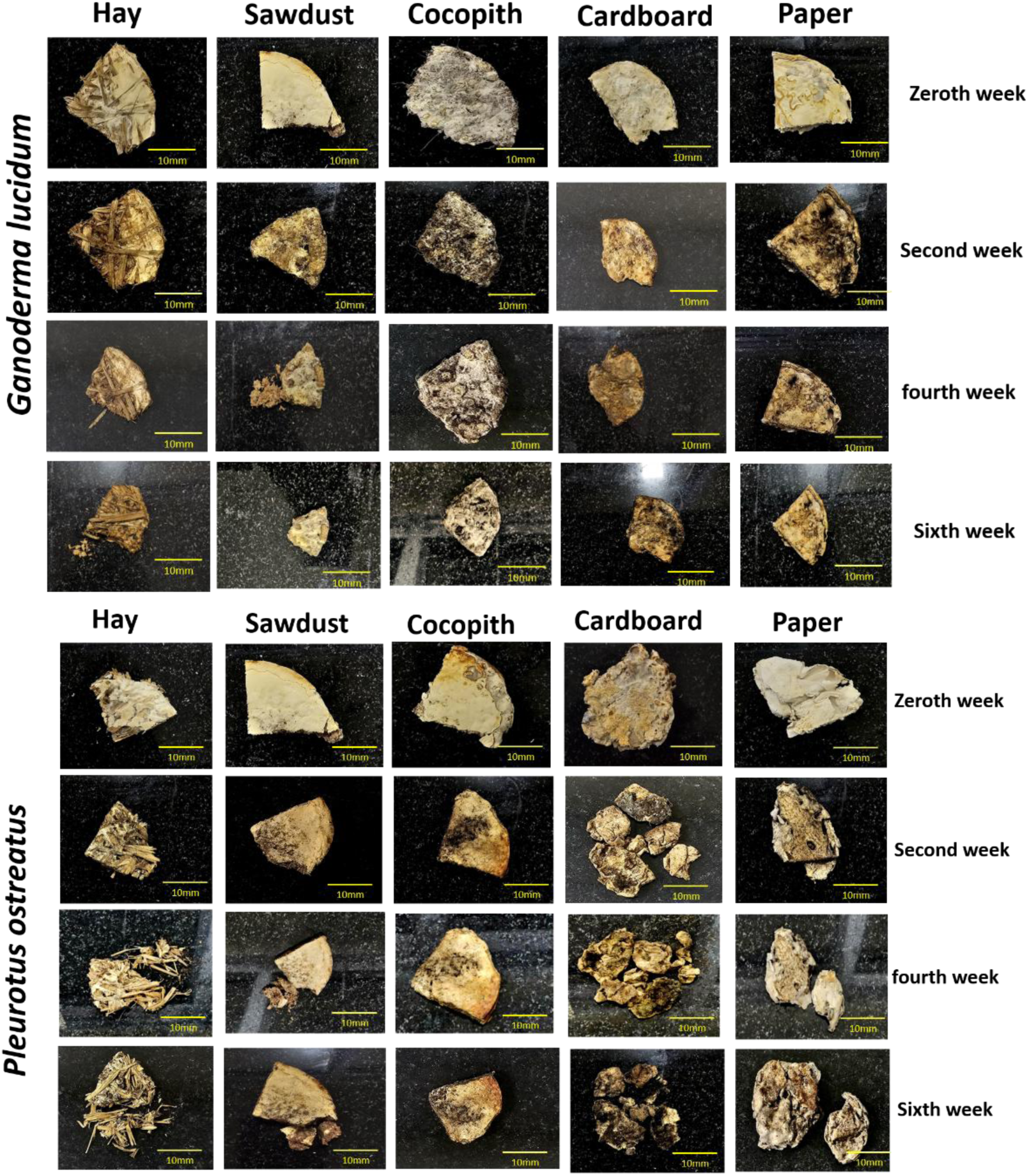
visual analysis of the *Ganoderma lucidum* and *Pleurotus ostreatus* samples before (top row) after second week (second row), fourth week (third row) and sixth week (fourth row)

### 3.6 Biodegradation test

Biodegradation experiments showed that all the samples started to disintegrate within 2 weeks’ time. The samples started to darken, loose its weight and reduce in size as observed in **Figure 9**. The initial darkening and loss of integrity observed in the samples suggest that the mycelium is losing its ability to hold the substrate together effectively. The mycelium which acted as a binder tend to lose its integrity and breakdown more with time from week 1 to week 6 (**Figure 9)**. This was also observed by (Van Wylick et al., 2022b). This breakdown can be influenced by environmental stressors such as moisture and temperature fluctuations. The graphs in **Figure. 10** depict that 80% weight loss was observed in cardboard *Pleurotus* sample within 6 weeks’ time followed by hay 48.8%, sawdust (46.30%), paper (38.6%) and cocopith (28.48%). Whereas sawdust sample of *Ganoderma lucidum* showed a 41.51% weight loss followed by hay (39.18%), paper (33.7%), cardboard (32.8%), and cocopith (12.19%). Humidity conditions (60-65%) can be a pivotal reason for this weight loss and disintegration of samples. This can accelerate the degradation of the samples by creating an environment that is conducive to microbial growth. Sawdust samples from *Ganoderma lucidum* had a notable weight loss (41.51%), suggesting that some substrates might be more vulnerable to enzymatic or microbial degradation than others (Zimele *et al*., 2020). How well these substrates withstand degradation can be influenced by their composition; for example, lignocellulosic materials, such as sawdust (49.40% of cellulose, 24.59% of hemicellulose, 26.8% lignin) and hay (35.8 % of cellulose, 21.5% of hemicellulose, 24.4% lignin) may deteriorate at a different rate than paper or cardboard (40-50 % of cellulose, 24% of hemicellulose, 18% lignin).

**Figure 10.**
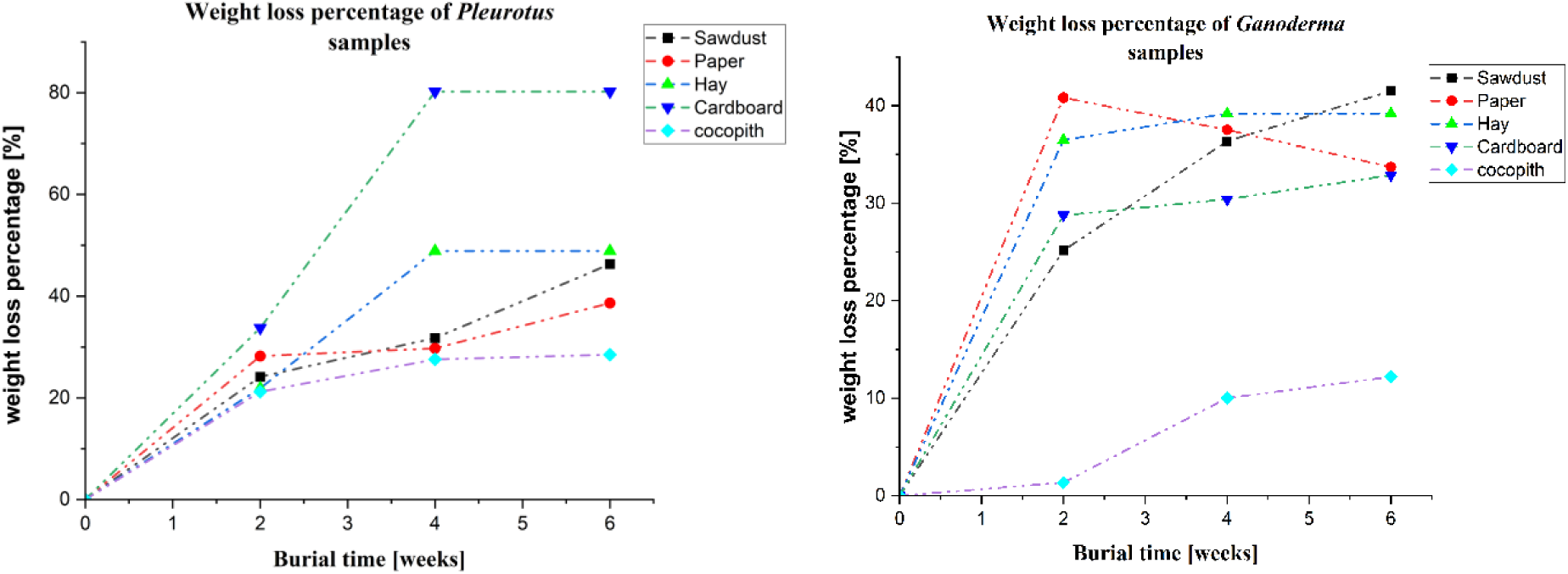
*W*eight loss percentage of *Ganoderma lucidum* and *Pleurotus ostreatus* samples after 6 weeks of burial in pot mix

## 4. Conclusions

The study aimed to evaluate the growth and material properties of mycelium biocomposites formed by *Ganoderma lucidum* and *Pleurotus ostreatus* grown on different substrates. The findings from the optical imaging and SEM analysis revealed that substrate type plays a significant role in mycelial growth, surface texture and density. Sawdust biocomposites of *Ganoderma* showed to the most favourable exhibiting the densest and smooth mycelial layers. The analysis also showed varying hyphal diameters across substrates. These structural differences were further linked to the mechanical and physical properties of the biocomposites.

The mycelium bio-composites’ modulus ranged from 1-3 MPa, meeting the Standard Specification for rigid EPS. The compression test results highlighted significant differences in the mechanical properties of biocomposites derived from various substrates and fungal strains. Cardboard emerged as the most effective substrate for *Ganoderma* yielding the highest compressive strength, while *Pleurotus* showed better results on sawdust. The findings underscore the critical role of substrate selection in optimizing growth conditions for each fungal strain, as the accumulation of chitin in the fungal cell walls contributes to enhanced mechanical resistance. Lighter substrates had low compressive strengths.

Water absorption tests revealed the correlation between hyphal structure and water retention capacity. It was also dependent on the porous nature of the substrates. In contrast to substrates like paper and cardboard, high porous substrates like hay absorbed a larger amount of water. Water contact angle tests confirmed that biocomposites with denser and smoother surfaces particularly *Pleurotus* sawdust and *Ganoderma* cardboard samples, had higher hydrophobicity. This suggests that the surface properties of the mycelium are key to understanding its potential applications in areas requiring moisture resistance.

The microstructural variations observed in mycelium-based composites formed from different substrates and fungus species, such as differences in hyphal diameter, surface porosity, and network formation, indicate that the choice of substrate and fungus species significantly influences the final properties and characteristics. Understanding these properties is essential for optimizing the performance and applications of these eco-friendly materials in various application. Future studies aimed at further tuning the properties by varying the parameters of the substrate, species of fungi and growth conditions, will lead towards biocomposites that can closely mimic the properties of packaging materials and still be biodegradable.

## Acknowledgments

The authors would like to thank Prof. Arockiarajan Arunachalakasi for giving access to the compression testing equipment. This project was carried with support from the Department of Applied Mechanics and Biomedical Engineering, NFIG grant of IIT Madras and Ministry of Education, India.

## Author Contributions

L.K. conceptualized and designed study. S.R.B has performed the experiments and did the analysis. V.S carried out the compression tests and helped in the corresponding data analysis. S.R.B and L.K wrote and edited the manuscript.

